# The Brain Dynamics Toolbox for Matlab

**DOI:** 10.1101/219329

**Authors:** Stewart Heitmann, Matthew J Aburn, Michael Breakspear

**Affiliations:** QIMR Berghofer Medical Research Institute 300 Herston Road, Herston QLD 4006, Australia

**Keywords:** initial-value problems, differential equations, numerical integration, visualization, brain dynamics

## Abstract

Nonlinear dynamical systems are increasingly informing both theoretical and empirical branches of neuroscience. The Brain Dynamics Toolbox provides an interactive simulation platform for exploring such systems in MATLAB. It supports the major classes of differential equations that arise in computational neuroscience: Ordinary Differential Equations, Delay Differential Equations and Stochastic Differential Equations. The design of the graphical interface fosters intuitive exploration of the dynamics while still supporting scripted parameter explorations and large-scale simulations. Although the toolbox is intended for dynamical models in computational neuroscience, it can be applied to dynamical systems from any domain.

## 1. Introduction

Computational neuroscience relies heavily on numerical methods for simulating non-linear models of brain dynamics. Software toolkits are the manifestation of those endeavors. Each one represents an attempt to balance mathematical flexibility with computational convenience. Toolkits such as GENESIS [1], NEURON [2] and BRIAN[3] provide convenient methods to simulate conductance-based models of single neurons and networks thereof. The Virtual Brain [4] scales up that approach to the macroscopic dynamics of the whole brain by combining neural field models [5] with anatomical connectivity datasets [6]. Mathematical toolkits such as AUTO [7], XPPAUT [8], MATCONT [9], PyDSTool [10] and CoCo [11] are useful for analyzing non-linear dynamics but assume advanced mathematical theory.

## 2. Problems and Background

In our experience, the existing computational toolkits often present technical barriers to broader audiences in cognitive neuroscience, systems neuroscience and neuroimaging. For example, GENESIS [1], NEURON [2], BRIAN [3] and XPPAUT [8] each use idiosyncratic languages for defining the differential equations. The Virtual Brain [4], CoCo [11] and PyDSTool [10] use conventional programming languages (Python and MATLAB) but assume advanced object-oriented programming techniques that broader audiences often find confusing. Of all of the existing toolkits, only XPPAUT [8] and the Virtual Brain [4] are capable of supporting Ordinary Differential Equations (ODEs), Delay Differential Equations (DDEs) and Stochastic Differential Equations (SDEs). Our Brain Dynamics Toolbox aims to bridge these technical barriers by allowing those with diverse backgrounds to explore neuronal dynamics through phase space analysis, time series exploration and other methods with minimal programming burden. A custom system of ODEs, DDEs or SDEs can typically be implemented in fewer than 100 lines of standard MATLAB code. Object-oriented programming techniques are not required. Once the model is implemented, it can be run interactively in the graphical interface (Figure 1) where a variety of different plotting panels and numerical solvers can be applied with no additional programming effort. The internal states of the graphical interface are accessible to the user’s workspace so that parameter sweeps can be semi-automated in the command window with simple for-loop statements. Additional command-line tools are also provided for scripting fully-automated simulations in batch mode. Such scripts may be called from third-party MATLAB applications and vice versa. Large-scale simulations can be scripted to run in parallel using the MATLAB Parallel Computing Toolbox or the MATLAB Distributed Computing Server. Unfortunately the toolbox does not run on *Octave* [12] because of incompatibilities in the graphical interface class libraries.

## 3. Software Framework

The toolbox operates on user-defined systems of ODEs, DDEs and SDEs. The details differ slightly for each type of differential equation but the overall approach is the same. For an ODE,

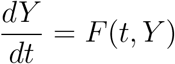

the right-hand side of the equation is implemented as a matlab function of the form `dYdt=F(t,Y)`. The toolbox takes a handle to that function and passes it to the relevant solver routine on the user’s behalf. The solver repeatedly calls `F(t,Y)` in the process of computing the evolution of *Y*(*t*) from a given set of initial conditions. The toolbox uses the same approach as the standard MATLAB solvers (e.g. `ode45`) except that it also manages the input parameters and plots the solver output. To do so, it requires the names and values of the system parameters and state variables. Those details (and more) are passed to the toolbox via a special data structure that we call a *system structure*. It encapsulates everything needed to simulate a user-defined model. Once a system structure has been constructed, it can be shared with other toolbox users.

**Figure 1.**
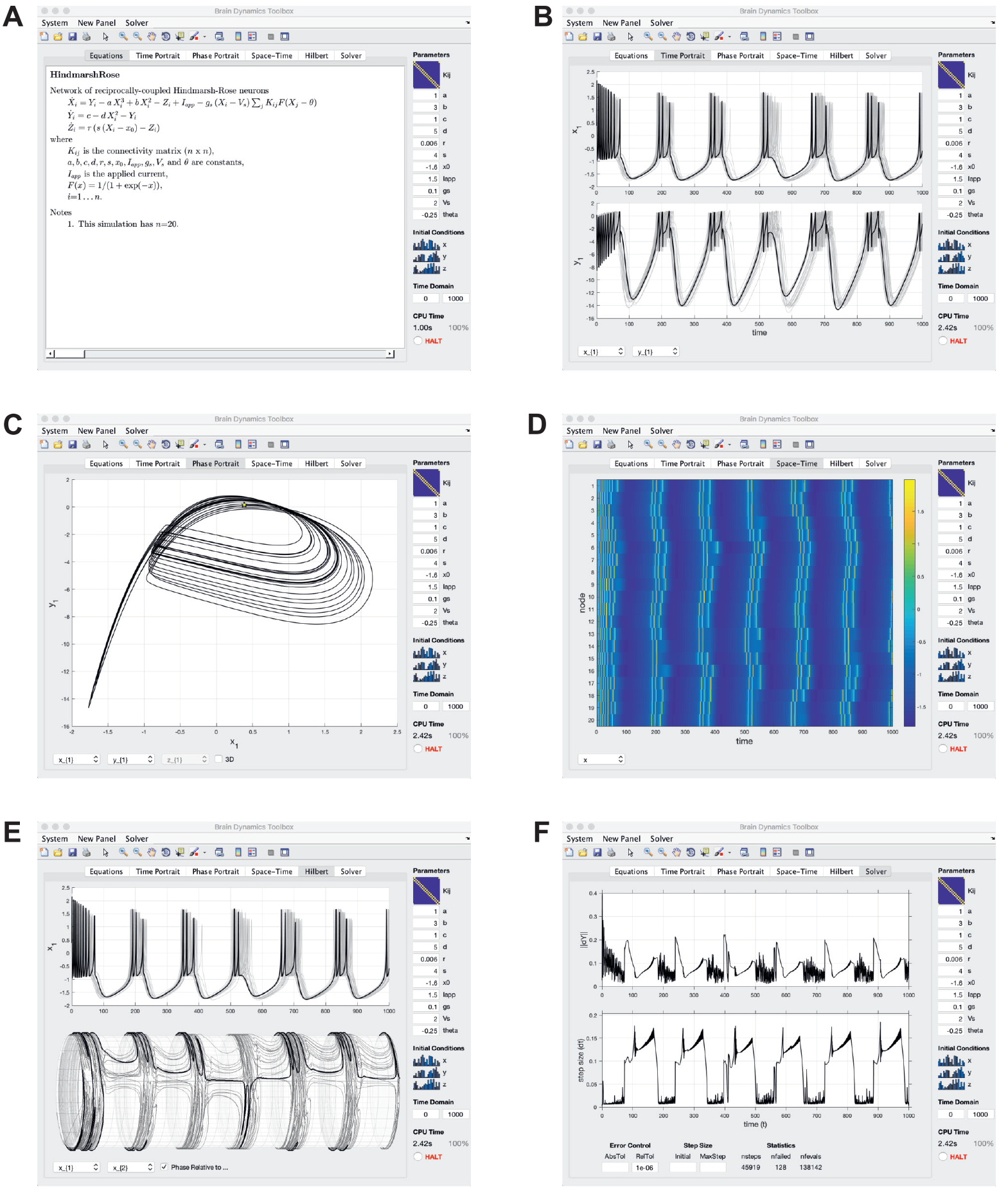
Screenshots of selected display panels in the graphical interface as it simulates a network of *n*=20 Hindmarsh-Rose [13] neurons. The parameters of the model appear in the control panel on the right-hand side of the application window. The solution is automatically recomputed each time any of those controls are altered. Individual controls can be scalar, vector or matrix values thereby accommodating arbitrarily large parameter sets. **A** Mathematical equations rendered with LaTeX. **B** Time portraits. **C** Phase portrait. **D** Space-time portrait. **E** Hilbert transform. **F** Solver step sizes.

### 3.1. Software Architecture and Functionality

The hub-and-spoke software architecture (Figure 2) allows arbitrary combinations of solver routines and display panels to be applied to any model. The modular design also allows new solver routines and display panel classes to be added to the toolbox incrementally. The list of numerical solver routines and graphical panels that the toolbox supports continues to grow rapidly. The current version (2017c) supports the standard ODE solvers (`ode45, ode23, ode113, ode15s, ode23s`) and DDE solver (`dde23`) that are shipped with MATLAB. As well as a fixed-step ODE solver (`odeEul`) and two SDE solvers (`sdeEM, sdeSH`) that are specific to the Brain Dynamics Toolbox. The two SDE solvers are specialized for stochastic equations that use Itô calculus and Stratonovich calculus respectively.

The display panels can be used to visualize the dynamics, compute metrics from the time-series, or transform them into new time-series. The toolbox currently includes display panels for rendering mathematical equations, time plots, phase portraits, space-time plots, computing linear correlations, Hilbert transforms, surrogate data transforms and inspecting the individual steps taken by the solvers. The panel outputs are themselves accessible to the user’s workspace as read-only variables. New panels can be added to the toolbox at any time and we encourage advanced users to write custom panels for their own projects although that level of graphical interface development does involve object-oriented programming.

**Figure 2:**
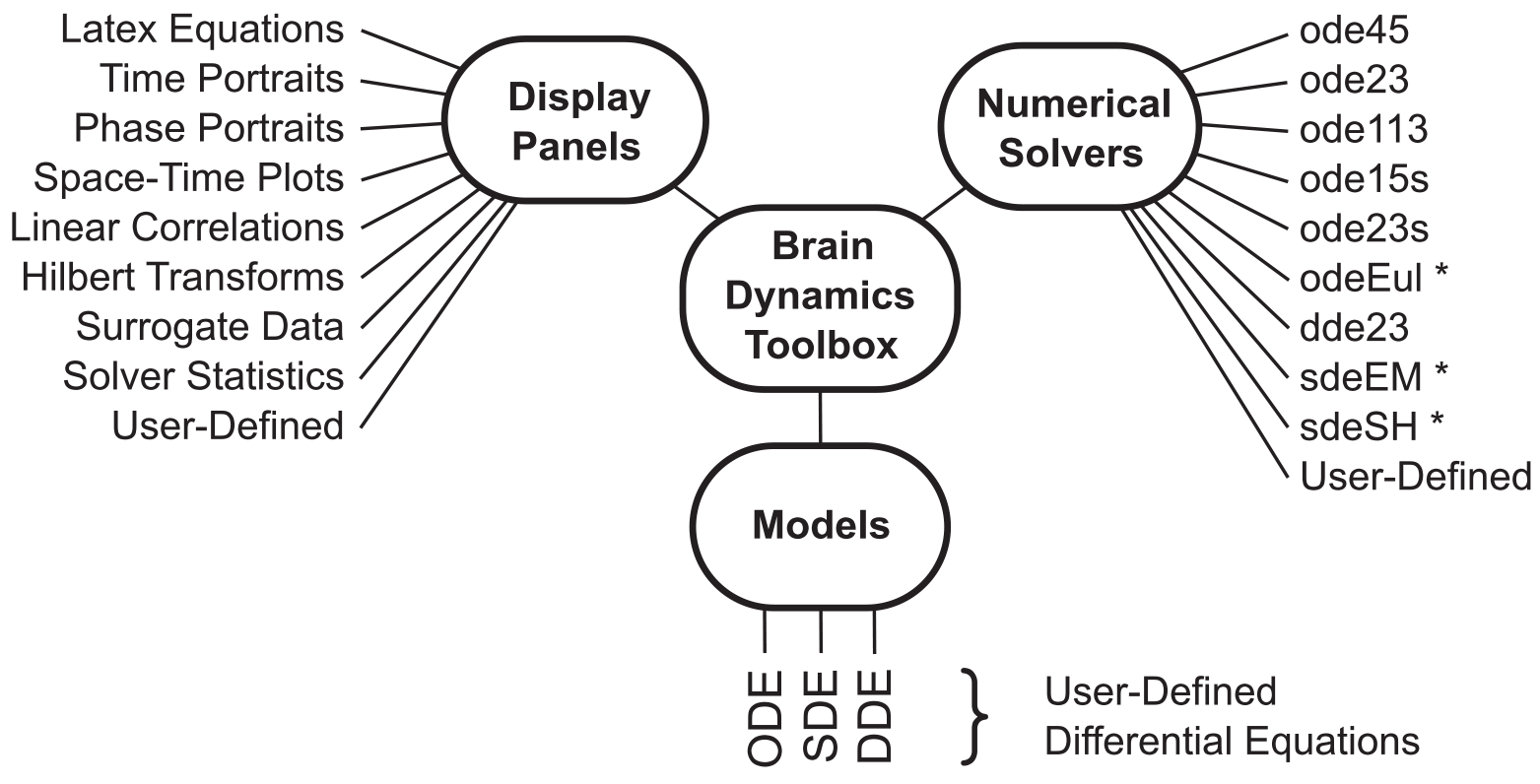
The hub-and-spoke software architecture of the Brain Dynamics Toolbox. Numerical solvers marked with an asterisk are unique to the toolbox.

## 4. Illustrative Example

We demonstrate the implementation of a network of recurrently-connected Hindmarsh-Rose [13] neurons,

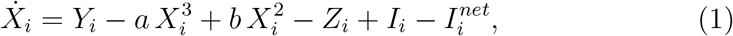

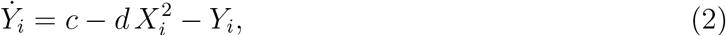

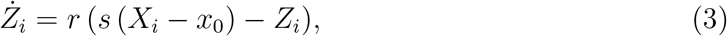

where *X*_*i*_ is the membrane potential of the *i*^*th*^ neuron, *Y*_*i*_ is the conductance of that neuron’s excitatory ion channels, and *Z*_*i*_ is the conductance of its inhibitory ion channels. Each neuron in the network is driven by a locally applied current *I*_*i*_ and a network current 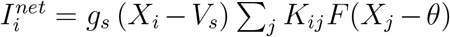 that represents the synaptic bombardment from other neurons. The sigmoidal function *F*(*x*)=1/(1 + exp(−*x*)) transforms that synaptic bombardment to an equivalent ionic current. The connectivity matrix *K*_*ij*_ defines the weightings of the synaptic connections between neurons. All other parameters in the model are scalar constants. The model is a typical example of a neuronal network as a system of coupled ODEs.

### 4.1. Defining the equations

We define the right-hand side of equations (1-3) as MATLAB function of the form `dY=F(t,Y,…)` where the vector `Y` contains the instantaneous values of [*X*(*t*), *Y*(*t*), *Z*(*t*)] at time *t*. The ellipses denote model-specific parameters.

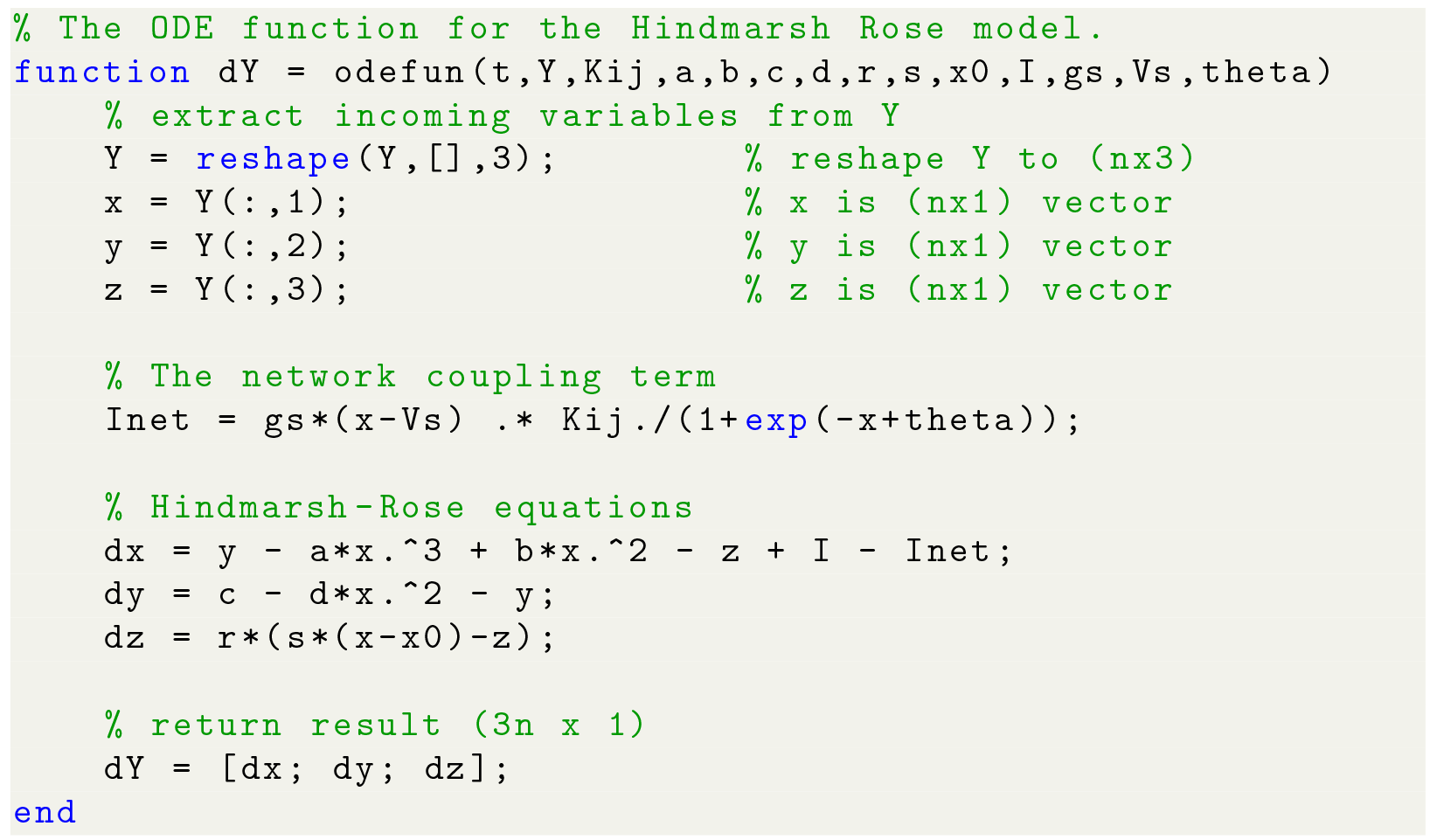

It is no coincidence that the form of this function is identical to that required by the standard ODE solvers, since the toolbox applies those same solvers to this function. In order to do so, it requires the names and initial values of the model’s parameters and state variables to also be defined. That is the purpose of the model’s system structure.

### 4.2. Defining the system structure

The system structure (named `sys` by convention) encapsulates the function handles and parameter settings that the toolbox needs to pass the user-defined ODE function to the solver and plot the solution that is returned. The most important fields of the structure are the handle to user-defined function (`sys.odefun`), the names and initial values of the system variables (`sys.vardef`) and the names and values of the system parameters (`sys.pardef`). Once a system structure has been constructed, it can be saved to a mat file and used by the toolbox as is. Nonetheless it is common practice to provide a helper function that constructs a new system structure for a particular configuration of the model. In this example, the configuration of the system variables depends on the size of the network connectivity matrix, `Kij`.

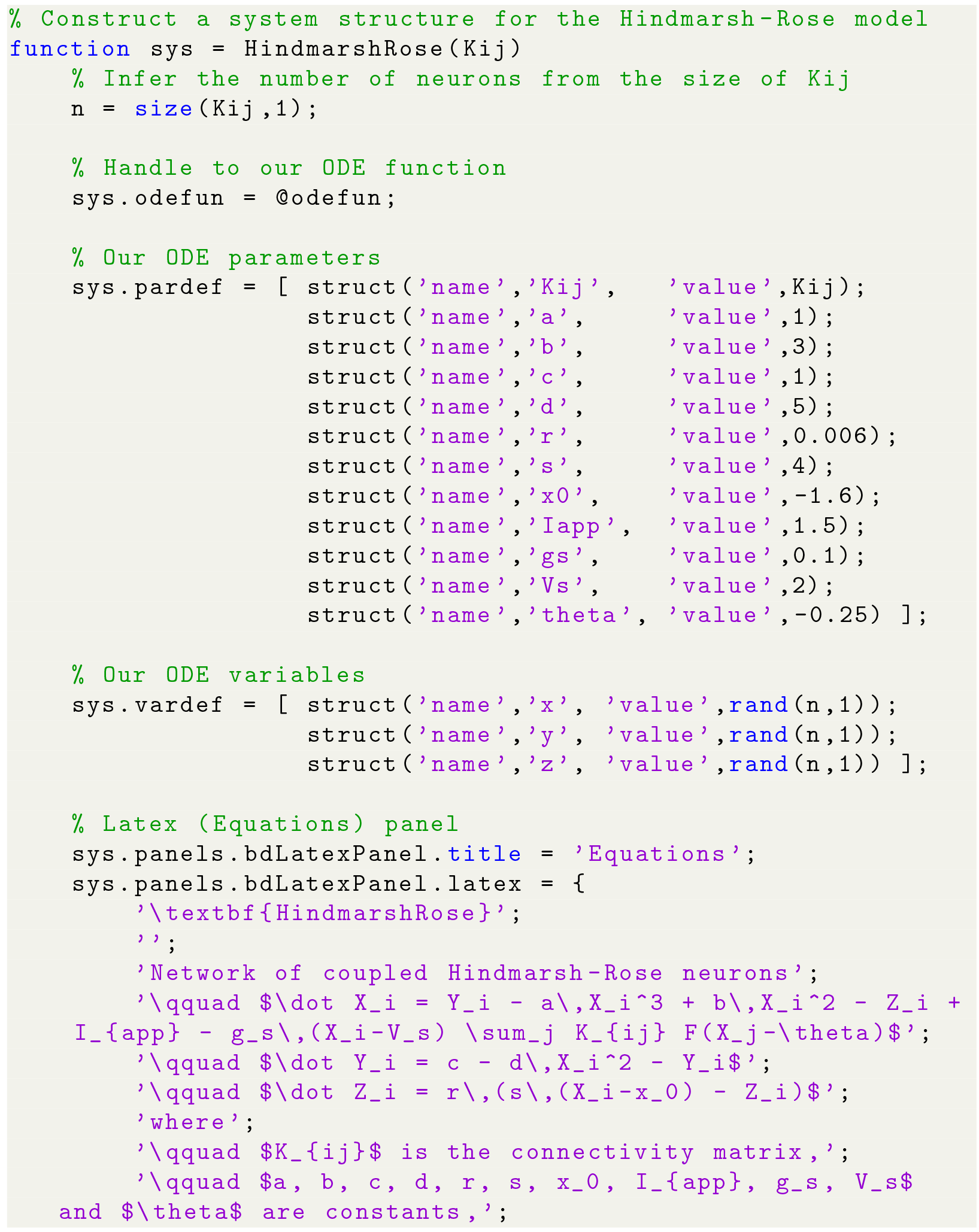

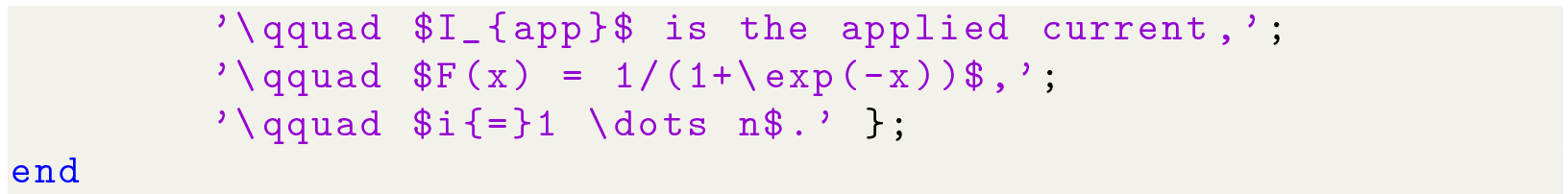

The order of the parameter definitions in the `pardef` field must match that of the `odefun` function. Likewise for the system variables in the `vardef` field.

The final part of the helper function (lines 28-42) defines the model-specific strings for rendering the mathematical equations in the LaTeX display panel. Those LaTeX strings are important for documenting the model in the graphical interface but they play no part in the simulation itself.

### 4.3. Running the model

The model is run by loading an instance of the system structure into the toolbox graphical user interface, which is called `bdGUI`.

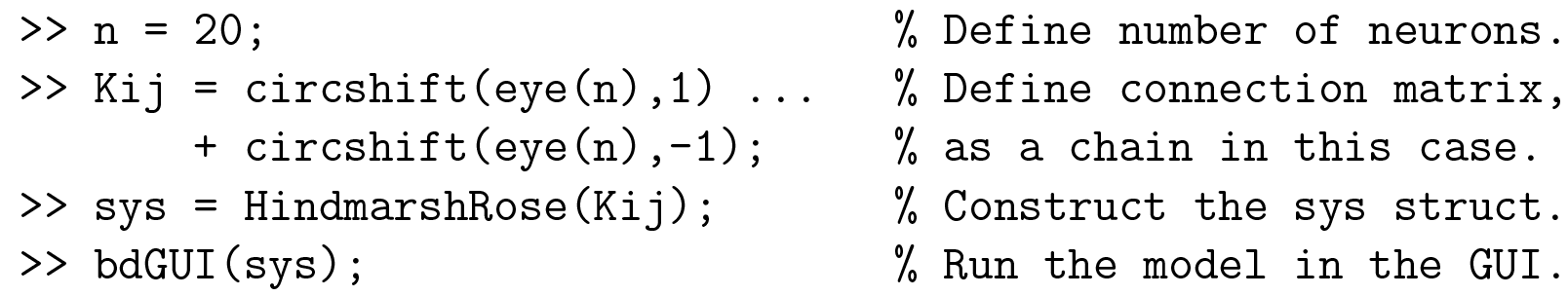

The graphical interface (Figure 1) allows the solution to be visualized with any number of display panels, all of which are updated concurrently. The solution is automatically recomputed whenever any of the graphical controls are adjusted; including the system parameters, the initial conditions of the state variables, the time domain of the simulation and the solver options.

### 4.4. Controlling the model

The `bdGUI` application returns a handle to itself which can be used to control the simulation from the MATLAB command window.

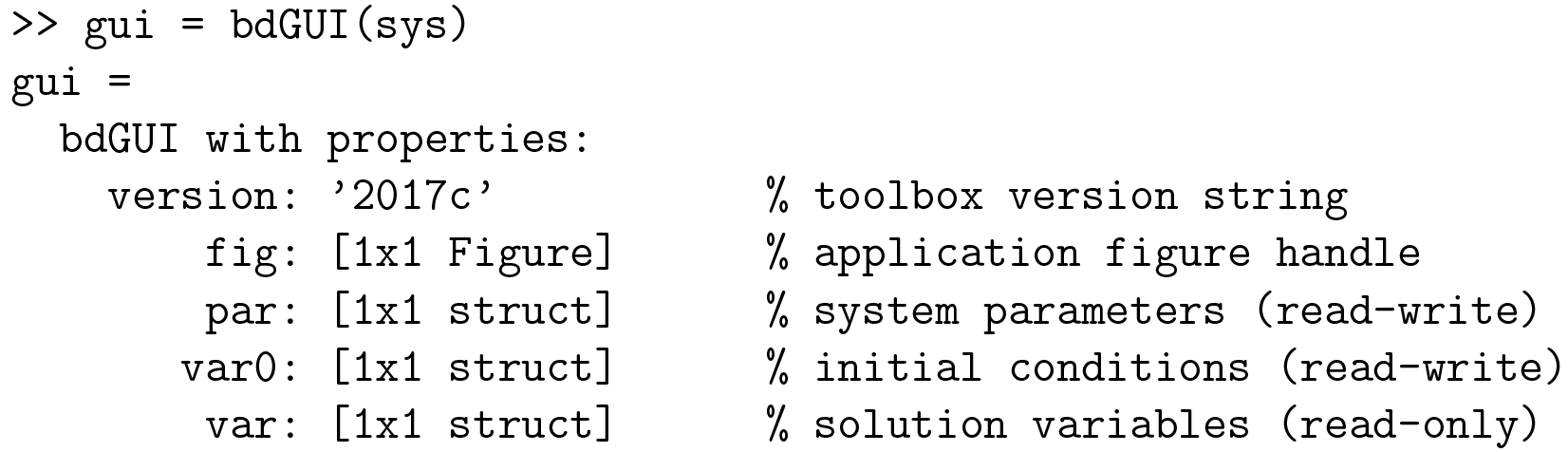

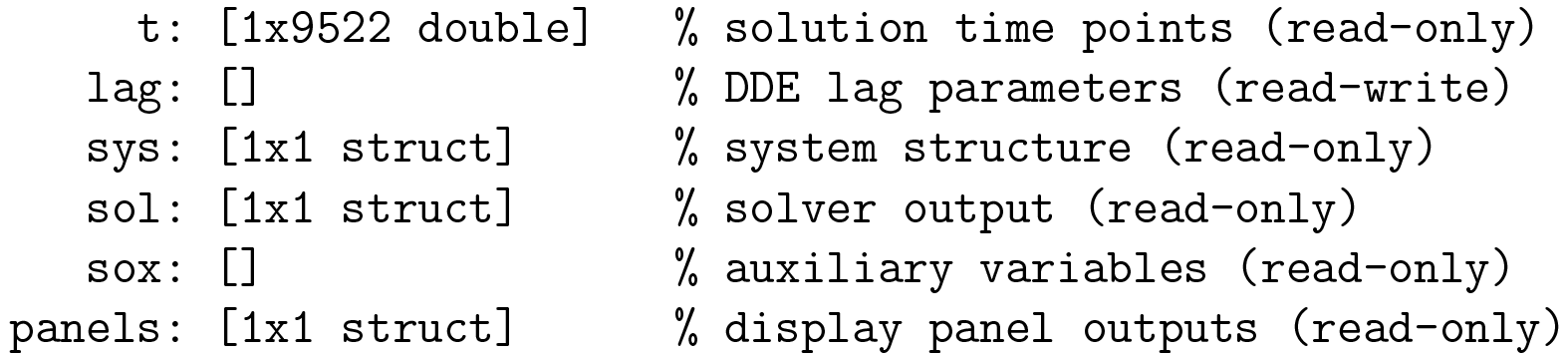

The parameters of the model are all accessible by name via the `gui.par` structure. Likewise, the computed solution variables are accessible by name via the `gui.var` structure and also in the native format returned by the solver via the `gui.sol` structure. Parameter values written into the `gui.par` handle are immediately applied to the graphical user interface, and vice versa. Hence it is possible to use workspace commands to orchestrate parameter sweeps in the graphical user interface. For example, the workspace command

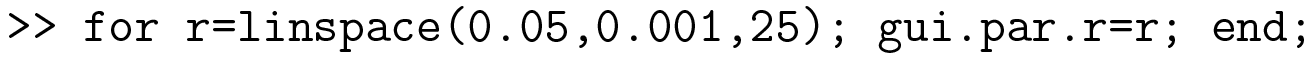

sweeps the *r* parameter (time constant of inhibition) from 0.05 to 0.001 in 25 increments. The graphical interface automatically recomputes the solution every time that `gui.par.r` is assigned a new value in the loop. The result is an animated sequence of simulations where bursting phenomenon is observed for *r* ≲ 0.01.

### 4.5. Scripting the model

The toolbox also provides a small suite of command-line tools for running models without invoking the graphical interface. Of these, the most notable commands are `bdSolve(sys,tspan)` which runs the solver on a given model for a given time span; and `bdEval(sol,t)` which interpolates the solution for a given set of time points.

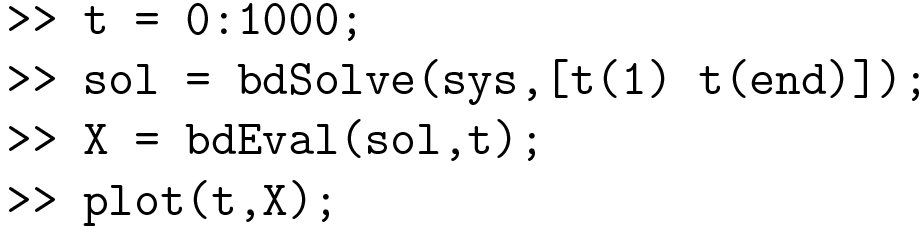

The `bdEval` function is equivalent to the MATLAB `deval` function except that it also works for solution structures (`sol`) returned by third-party solvers.

## 5. Conclusions

The Brain Dynamics Toolbox provides researchers with an interactive graphical tool for exploring user-defined dynamical systems without the burden of programming bespoke graphical applications. The graphical interface imposes no limit the size of the model nor the number of parameters involved. System parameters and variables can range in size from simple scalar values to large-scale vectors or matrices without loss of generality. The design also imposes no barrier to scripting large-scale simulations and parameter surveys. The toolbox is aimed at students, engineers and researchers in computational neuroscience but it can also be applied to general problems in dynamical systems. It is supported with an extensive user manual [14] that provides detailed instructions for implementing new systems of ODEs, DDEs and SDEs. Once a new model is implemented, it can be readily shared with other toolbox users. The toolbox thus serves as a hub for sharing models as much as it serves as a tool for simulating them.

## Acknowledgements

MATLAB^®^ is a registered trademark of The Mathworks, Inc., 3 Apple Hill Drive, Natick, MA 01760-2098 USA, 508-647-7000, Fax 508-647-7001, www.mathworks.com

## Required Metadata

### Current executable software version

**Table 1.** Software metadata (optional)

| Nr. | (executable) Software metadata description | Please fill in this column |
| --- | --- | --- |
| S1 | Current software version | 2017c |
| S2 | Permanent link to executables of this version | https://github.com/breakspear/bdtoolkit/releases/tag/bdtoolkit-2017c |
| S3 | Legal Software License | BSD 2-clause |
| S4 | Computing platform/Operating System | Matlab 2014b or newer |
| S5 | Installation requirements & dependencies | Signal Processing Toolbox (optional). Statistics and Machine Learning Toolbox (optional). |
| S6 | If available, link to user manual - if formally published include a reference to the publication in the reference list | http://www.bdtoolbox.org |
| S7 | Support email for questions |  |

### Current code version

**Table 2.** Code metadata (mandatory)

| **Nr.** | **Code metadata description** | **Please fill in this column** |
| --- | --- | --- |
| C1 | Current code version | 2017c |
| C2 | Permanent link to code/repository used of this code version | https://github.com/breakspear/bdtoolkit/releases/tag/bdtoolkit-2017c |
| C3 | Legal Code License | BSD 2-clause |
| C4 | Code versioning system used | git |
| C5 | Software code languages, tools, and services used | Matlab 2014b or newer |
| C6 | Compilation requirements, operating environments & dependencies | Signal Processing Toolbox (optional). Statistics and Machine Learning Toolbox (optional). |
| C7 | If available Link to developer documentation/manual | http://www.bdtoolbox.org |
| C8 | Support email for questions |  |

